# Absence of entourage: Terpenoids commonly found in *Cannabis sativa* do not modulate the functional activity of Δ9-THC at human CB1and CB2 receptors

**DOI:** 10.1101/569079

**Authors:** Marina Santiago, Shivani Sachdev, Jonathon C Arnold, Iain S McGregor, Mark Connor

**Author notes:** Corresponding author: Marina Santiago. Some of these data were presented at The Australasian Society of Clinical and Experimental Pharmacologists and Toxicologists Conference in Melbourne, Australia (2016).

## Abstract

**Introduction:** Compounds present in *Cannabis sativa* such as phytocannabinoids and terpenoids, may act in concert to elicit therapeutic effects. Cannabinoids such as Δ^9^-tetrahydrocannabinol (Δ^9^-THC) directly activate cannabinoid receptor 1 (CB_1_) and cannabinoid receptor 2 (CB_2_), however, it is not known if terpenoids present in *Cannabis* also affect cannabinoid receptor signalling. Therefore, we examined 6 common terpenoids alone, and in combination with cannabinoid receptor agonists, on CB_1_ and CB_2_ signalling *in vitro*.

**Materials and Methods:** Potassium channel activity in AtT20 FlpIn cells transfected with human CB_1_ or CB_2_ receptors was measured in real-time using FLIPR^®^ membrane potential dye in a FlexStation 3 plate reader. Terpenoids were tested individually and in combination for periods up to 30 minutes. Endogenous somatostatin receptors served as a control for direct effects of drugs on potassium channels.

**Results:** α-Pinene, β-pinene, β-caryophyllene, linalool, limonene and β-myrcene (up to 30-100 µM) did not change membrane potential in AtT20 cells expressing CB1 or CB2, or affect the response to a maximally effective concentration of the synthetic cannabinoid CP55,940. The presence of individual or a combination of terpenoids did not affect the hyperpolarization produced by Δ^9^-THC (10µM): (CB1: control, 59±7%; with terpenoids (10 µM each) 55±4%; CB2: Δ^9^-THC 16±5%, with terpenoids (10 µM each) 17±4%). To investigate possible effect on desensitization of CB1 responses, all six terpenoids were added together with Δ^9^-THC and signalling measured continuously over 30 min. Terpenoids did not affect desensitization, after 30 minutes the control hyperpolarization recovered by 63±6%, in the presence of the terpenoids recovery was 61±5%.

**Discussion:** None of the six of the most common terpenoids in *Cannabis* directly activated CB1 or CB2, or modulated the signalling of the phytocannabinoid agonist Δ^9^-THC. These results suggest that if a phytocannabinoid-terpenoid entourage effect exists, it is not at the CB1 or CB2 receptor level. It remains possible that terpenoids activate CB1 and CB2 signalling pathways that do not involve potassium channels, however, it seems more likely that they may act at different molecular target(s) in the neuronal circuits important for the behavioural effect of *Cannabis*.

## Introduction

An enduring notion in the medicinal *Cannabis* and cannabinoid field is that of entourage: the idea that use of the whole plant may exert substantially greater effects than the sum of its individual parts.^1^ Entourage is usually construed as a positive attribute, with the assumption that superior therapeutic actions, or a more favourable “high”, will be obtained from consuming the whole *Cannabis* plant rather than individual components such as Δ^9^-tetrahydrocannabinol (THC). Somewhat surprisingly, the evidence for this widely cited notion is relatively sparse.

*Cannabis* contains almost 150 phytocannabinoids, the most common of which are THC and cannabidiol (CBD), together with their acid precursors THCA and CBDA^2^. *Cannabis* also contains a large number of monoterpene and sesquiterpene compounds (together called terpenoids), the most common of which include α-pinene, β-pinene, linalool, limonene and β-myrcene (monoterpenes) and β-caryophyllene and caryophyllene oxide (sesquiterpenes).^3^ Terpenoids are volatile compounds that are synthesised alongside phytocannabinoids mainly in the trichomes of the cannabis plant, and provide cannabis with its distinctive aroma and flavour.^4^ Terpenoids are often lost if the extraction process involves heating.^5^

The entourage concept applied to cannabis can encompass the potential for both cannabinoid-cannabinoid and cannabinoid-terpenoid interactions. With regard to the former, Δ^9^-THC-CBD synergy in producing analgesia was reported in an animal model of neuropathic pain^6^ while in humans, CBD has been proposed to ameliorate some of the adverse psychotomimetic and anxiogenic effects of Δ^9^-THC.^7, 8^ This claim is controversial, however, with a number of contrary findings ^9, 10^ CBD may modulate Δ^9^-THC effects at the receptor level acting as a CB_1_ negative allosteric modulator^11^, providing some biological plausibility to a modulatory interaction.

Scientific evidence for cannabinoid-terpenoid interactions is essentially absent and mostly comes from websites and dispensaries extolling the virtues of proprietary *Cannabis* chemical varieties, or chemovars.^12, 13^ However, some terpenoids do have intrinsic psychoactive and physiological effects, and modulatory effects on Δ^9^-THC actions is not farfetched.^1, 14^ For example, in studies with laboratory animals, limonene displayed anxiolytic effects, pinene increased gastrointestinal motility, linalool was sedative, anticonvulsant and anxiolytic, while myrcene produced sedation, analgesia and muscle relaxant effects (summarised in Russo and Marcu^14^). However, compelling evidence for cannabinoid-terpenoid interactions or synergy does not yet exist, and it is also worth noting the relatively low concentrations of terpenoids present in herbal cannabis. ^5, 13, 15, 16^

With so many bioactive components present in cannabis, the systematic, granular elucidation of possible entourage effects poses a substantial combinatorial puzzle and scientific challenge. As a preliminary approach to addressing this challenge, the present study examined whether the effects of Δ^9^-THC on its cognate cannabinoid receptors (CB_1_ and CB_2_) would be modified in the presence of terpenoids that are commonly found in cannabis, either alone or in combination. The demonstration of such a receptor-level entourage effect might lead to predictions regarding functional cannabinoid-terpenoid interaction effects that could be tested *in vivo*.

## Materials and Methods

### Cell culture

Experiments used mouse wild-type AtT20 FlpIn cells (AtT20-WT), or these cells stably transfected with human CB_1_ or CB_2_ receptors with 3x N-terminus haemagglutinin tags (AtT20-CB_1_ and AtT20-CB_2_respectively).^17^ Cells were cultivated in Dulbecco’s Modified Eagle’s Medium (DMEM; Sigma-Aldrich) supplemented with 10% foetal bovine serum (FBS; SIGMA/SAFC) and 100U penicillin/100µg streptomycin mL^−1^ (Gibco). Selection antibiotics were 80µg mL^−1^ Zeocin (Invivogen) for AtT20-WT or 80µg mL^−1^ hygromycin B Gold (Invivogen) for transfected cells.

Cells were grown in 75mm^2^ flasks at 37°C/5% CO_2_ and passaged when 80-90% confluent. Assays were carried out on cells up to 20 passages in culture.

### Potassium Channel Activity Measurements

Changes in membrane potential were measured using the FLIPR^®^ blue membrane potential dye (Molecular Devices) in a FlexStation 3, as outlined in Knapman 2013.^18^ Cells from a 90-100% confluent 75mm^2^ flask were resuspended in Leibovitz’s L-15 Medium (Gibco) supplemented with 1% FBS, 100U penicillin/100µg streptomycin mL^−1^ and glucose (15mM) and plated in 96 well black walled clear bottom microplates (Costar) in a volume of 90µL per well. Cells were incubated overnight in humidified ambient air at 37°C incubator. Membrane potential dye, used at 50% of the manufacturer recommended concentration, was resuspended in Hank’s Balanced Salt solution (HBSS) of composition (in mM): NaCl 145, HEPES 22, Na_2_HPO_4_ 0.338, NaHCO_3_ 4.17, KH_2_PO_4_ 0.441, MgSO_4_ 0.407, MgCl_2_ 0.493, CaCl_2_ 1.26, glucose 5.55 (pH 7.4, osmolarity 315±15). Dye was loaded into each well (90µL per well) and equilibrated at 37°C for at least 1 hour prior to assay. Fluorescence was measured every 2 seconds (λ excitation = 530nm, λ emission = 565nm, λ emission cut-off = 550nm). Assays were carried out at 37°C and drugs were automatically added in volumes of 20µL.

#### Determining the Effects of Terpenoids on Acute Hyperpolarization

Terpenoids were added after at least 60 seconds of baseline recording and incubated for 5 minutes before cannabinoid (CP55,940 or Δ^9^-THC) addition. In AtT20-WT cells, somatostatin (SST) was added instead of cannabinoid.

#### Determining the Effects of Terpenoids on Signalling Desensitization

Homologous desensitization was measured by simultaneously adding Δ^9^-THC with terpenoids after 120 seconds of baseline recording. Signalling desensitization was calculated as percentage decrease from peak Δ^9^-THC response after 30 minutes in drugs. SST (100nM) was added 30 minutes after Δ^9^-THC addition to examine the potential effects of prolonged cannabinoid receptor activation on native somatostatin receptors (heterologous desensitization). The SST response was compared between groups (with or without terpenoids).

#### Drug Dilution

All drugs (except SST) were made up in DMSO and stored as frozen stocks at a concentration of 10mM – 100mM. Terpenoid stock solution concentrations were 100mM, with exception of β-myrcene (30 mM) which was insoluble at 100mM. SST was dissolved in water. Fresh aliquots were used each day, with the drugs diluted in HBSS containing 0.1% bovine serum albumin (Sigma-Aldrich) immediately before the assay. The final concentration of DMSO in each well was 0.1 to 0.11%; this limited the maximum concentration of terpenoids able to be tested. A vehicle (HBSS plus solvent alone) well was included in each column of the 96 well plate and the changes in fluorescence produced by vehicle alone were subtracted before determining the maximum hyperpolarization after each drug exposure.

### Drugs and Reagents

Δ^9^-THC was from THCPharm (Frankfurt, Germany). Terpenoids were from Sigma-Aldrich; (+)-α-pinene, (+)-β-pinene, (-)-β-caryophyllene, (+/-)-linalool, (R)-(+)-limonene and β-myrcene. Somatostatin was from Auspep and CP55,940 from Cayman. Unless otherwise indicated, the other chemicals and reagents were from Sigma-Aldrich.

### Data Analysis

Each experiment was independently repeated at least 5 times, with 2 technical replicates in each determination. Data are expressed as a percentage change in the fluorescence compared with the pre-drug baseline (30s before drug addition), or as percentage of 1µM CP55,940 response. Graphs were plotted using Graphpad Prism 7.02, and scatter dot plots show means with standard error of the mean (SEM). Means were compared using unpaired Student’s *t*-test or no matching one-way ANOVA followed by correction for multiple comparisons (Dunnett); and null hypothesis was rejected if *p*-value was lower than 0.05 (*p* > 0.05 = not significant).

## Results

### Terpenoids in AtT20-WT cells

We first examined terpenoid action in non-transfected AtT20 cells. We used somatostatin (100nM) as a positive control because it hyperpolarizes AtT20-WT cells via activation of endogenous SST receptors (Fig. 1A and B).^18, 19^ Addition of α-pinene, β-pinene, β-caryophyllene, linalool, limonene (100μM) or β-myrcene (30μM) did not affect the membrane potential of AtT20-WT cells (Fig. 1C, open circles). The presence of terpenoids (100μM/30μM) had no effect on the subsequent somatostatin response (Fig. 1C).

**Fig. 1.**
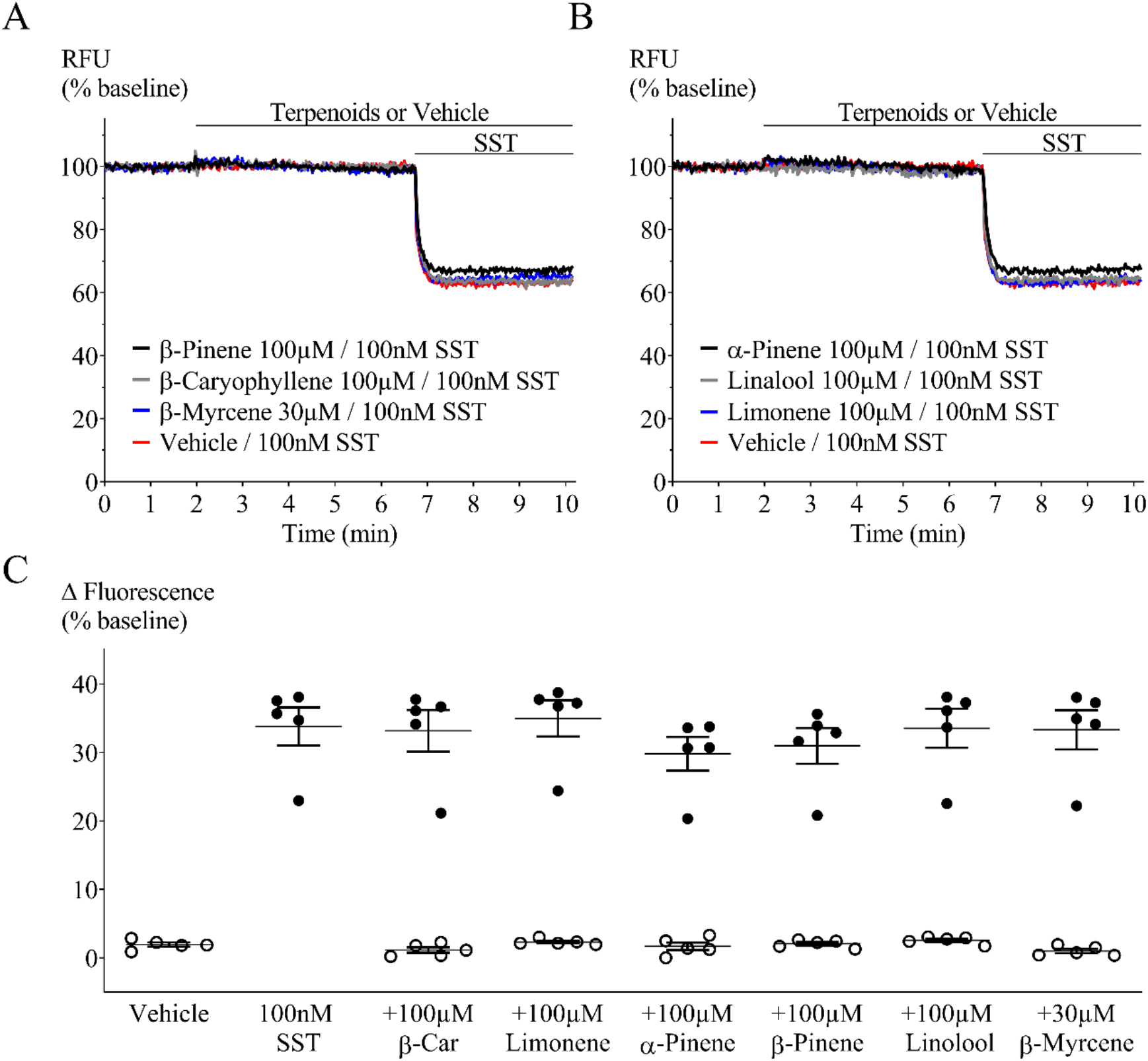
Terpenoid and somatostatin mediated fluorescence change in AtT20-WT. Representative traces showing change in fluorescence signal after terpenoid and somatostatin (SST, 100nM) application. A decrease in signal corresponds to membrane hyperpolarization. Addition of terpenoids (**A.** β-pinene, β-caryophyllene and β-myrcene; **B.** α-pinene, linalool and limonene) did not change baseline fluorescence, while somatostatin mediated a clear hyperpolarization. **C.** Percentage change of fluorescence from baseline after each terpenoid (open circle) and somatostatin (closed circles) application. Terpenoids were added at two minutes; five minutes before somatostatin. When compared to positive (SST) or negative (vehicle) controls, none of the terpenoids tested affected baseline membrane potential or peak somatostatin response. β-Car = β- caryophyllene. n = 5, SEM, one-way ANOVA *p* > 0.05. Drugs were added for the duration of the bar.

### Terpenoids in AtT20-CB_1_ and -CB_2_ cells

The absence of a terpenoid response in AtT20-WT cells enabled the study of their effect on membrane potential in AtT20 cells expressing human CB_1_ or CB_2_. We examined whether terpenoids (1nM – 100μM, β- myrcene 300pM - 30μM) hyperpolarised cells via these receptors and, in parallel, whether they affected a subsequent response to a maximally effective concentration of CP55,940 (1µM, Fig. 2).^17^ A summary of the fluorescence change after terpenoid addition to AtT20-CB_1_ cells is shown in Figure 3 (closed circles). No difference between vehicle and terpenoids was observed. Further, none of the terpenoids changed the membrane potential of cells expressing CB_2_ (Supplementary Fig. S1). The change in fluorescence produced by the subsequent addition of the non-selective cannabinoid agonist CP55,940 (1μM) was also unaffected in both AtT20-CB_1_ and -CB_2_ (Fig. 3 and Supplementary Fig. S1 – open circles).

**Fig. 2.**
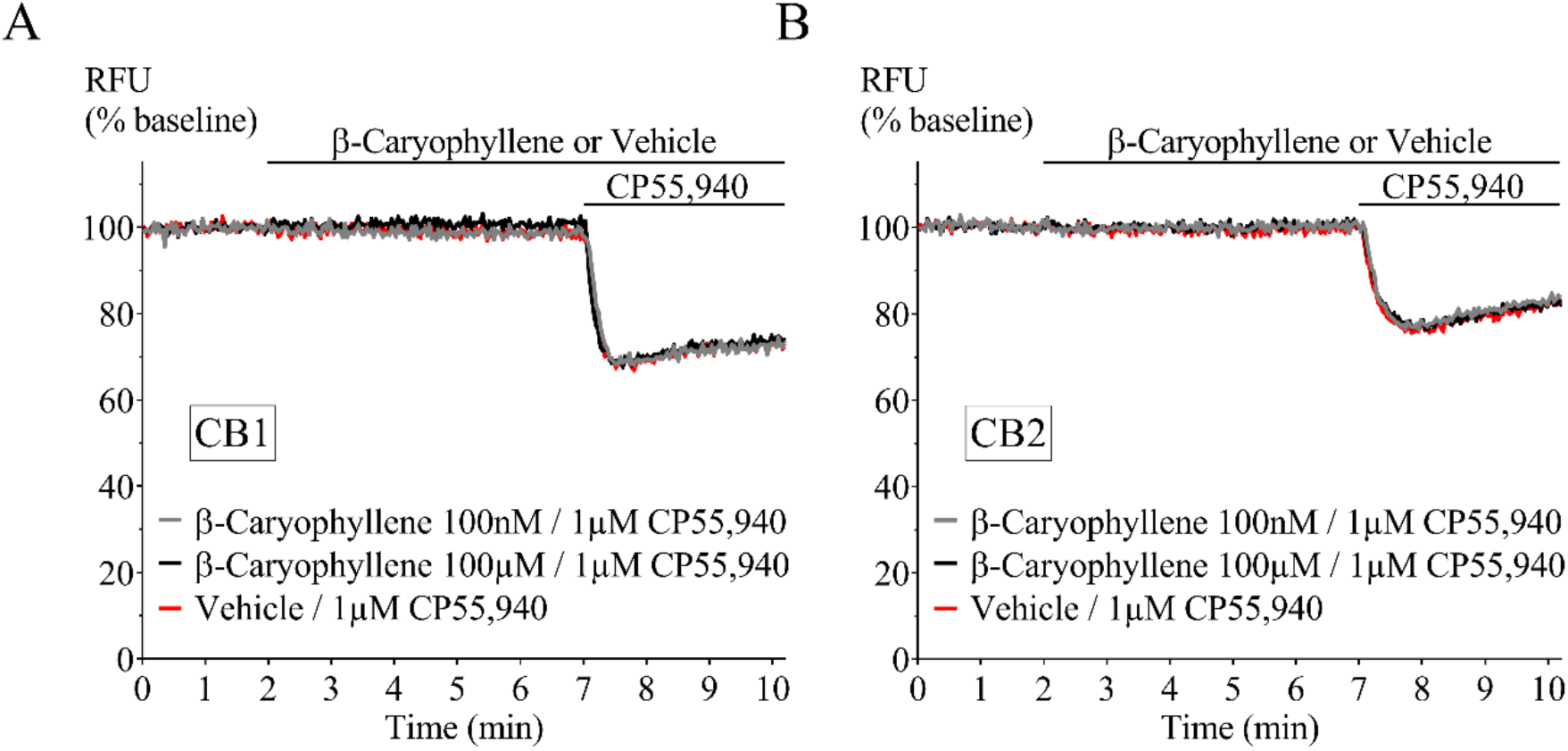
Representative traces of β-caryophyllene and CP55,940 in AtT20-CB1 and -CB2. Fluorescence was recorded for 10 minutes where β-caryophyllene (100nM and 100µM) was added at 2 minutes followed by incubation for 5 minutes, before 1µM CP55,940 application. β-caryophyllene did not hyperpolarize **A.** AtT20- CB1 and **B.** AtT20-CB2 cells, or affect the response to CP55,940 (1µM). Drugs were added for the duration of the bar.

**Fig. 3.**
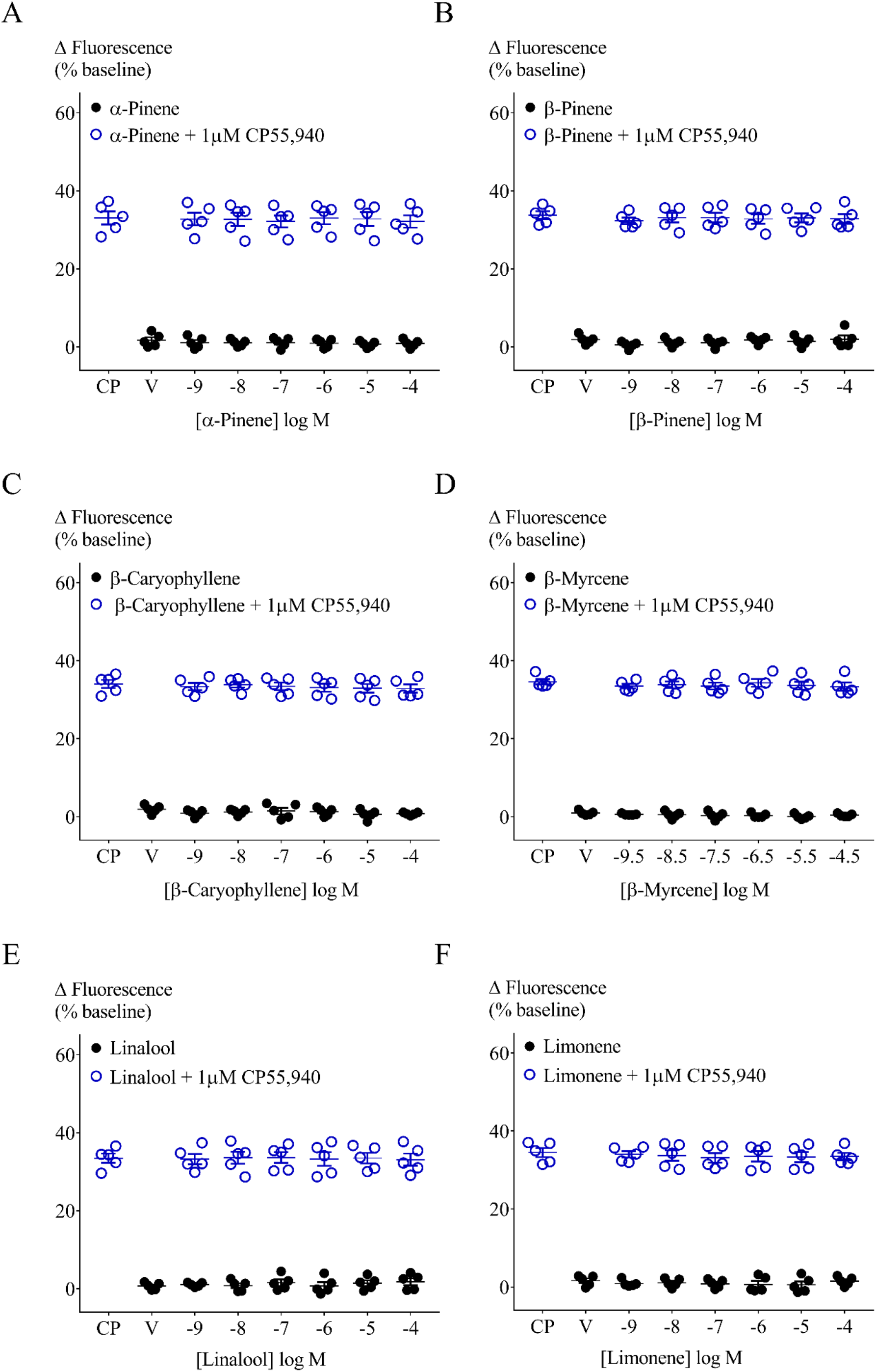
Effect of terpenoids at varying concentrations on AtT20-CB1 membrane potential and on 1μM CP55,940 induced hyperpolarization. Terpenoids (**A.** α-pinene, **B.** β-pinene, **C.** β-caryophyllene, **D.** β-myrcene, **E.** linalool and **F.** limonene) were added to AtT20-CB1 cells and incubated for 5 minutes. Maximum fluorescence changes were not different from negative control (closed circles, n=5, SEM, one-way ANOVA *p >*0.05). CP55,940 (1µM) addition to AtT20-CB1 cells induced fluorescence changes from 33.1 ±1.7% to 34.6±0.7%. Peak CP55,940 responses were not affected by the presence of terpenoids (open circles, n=5, SEM, one-way ANOVA *p >* 0.05). V = Vehicle.

**Fig. 4.**
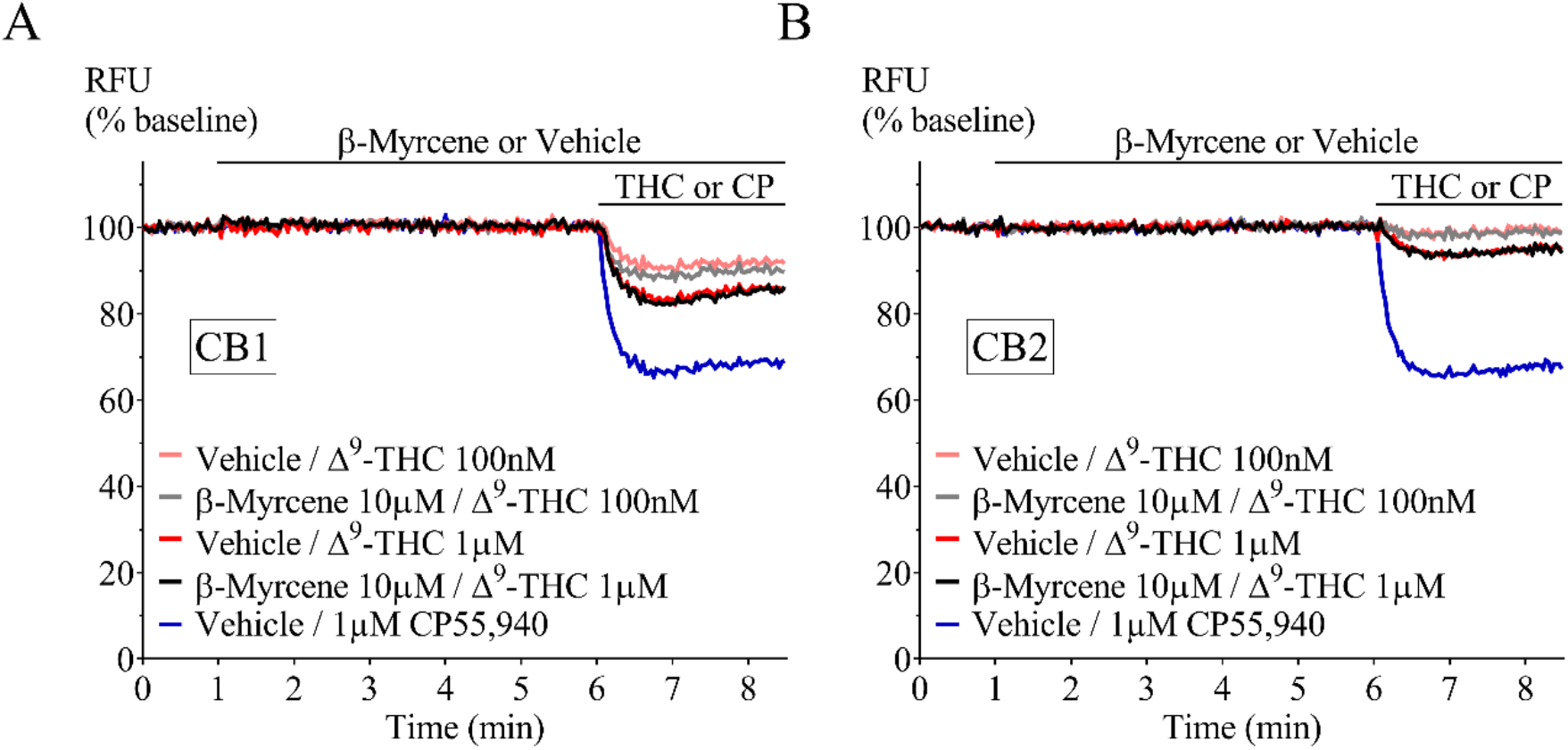
Representative traces of β-myrcene and Δ^9^-THC in AtT20-CB1 and AtT20-CB2. Fluorescence change mediated by two sub-maximal concentrations of Δ^9^-THC (100nM and 1µM) in the presence of β-myrcene (10µM). Terpenoid was added at 1 minutes and incubated for 5 minutes before Δ^9^-THC application. CP55,940 added as positive control. Drugs were added for the duration of the bar.

CP55,940 is a high efficacy agonist of both CB_1_ and CB_2_ receptors.^20^ However, in *Cannabis*, Δ^9^-THC is the principle cannabinoid agonist and it has a lower efficacy than CP55,940, which is apparent in the hyperpolarization assay as a lower maximal response.^20^ We next tested the effect of a low and high concentration of terpenoids (100nM and 10μM) on the hyperpolarization produced by three concentrations of Δ^9^-THC (100nM, 1μM and 10μM). After five minutes of individual terpenoid application, application of Δ^9^-THC produced fluorescence changes that were not significantly different from those produced by Δ^9^-THC alone in both AtT20-CB_1_ and -CB_2_ cells (10μM Δ^9^-THC Figs. 5 and 6, 100nM Δ^9^-THC Supplementary Figs. S2 and S3). To explore the possibility of an emergent entourage effect, we combined all six terpenoids (10μM each) and tested the effect of the mixture on the Δ^9^-THC-induced hyperpolarization. Similar to individually tested terpenoids, the effects of Δ^9^-THC were not changed by the mixture (Fig. 7).

**Fig. 5.**
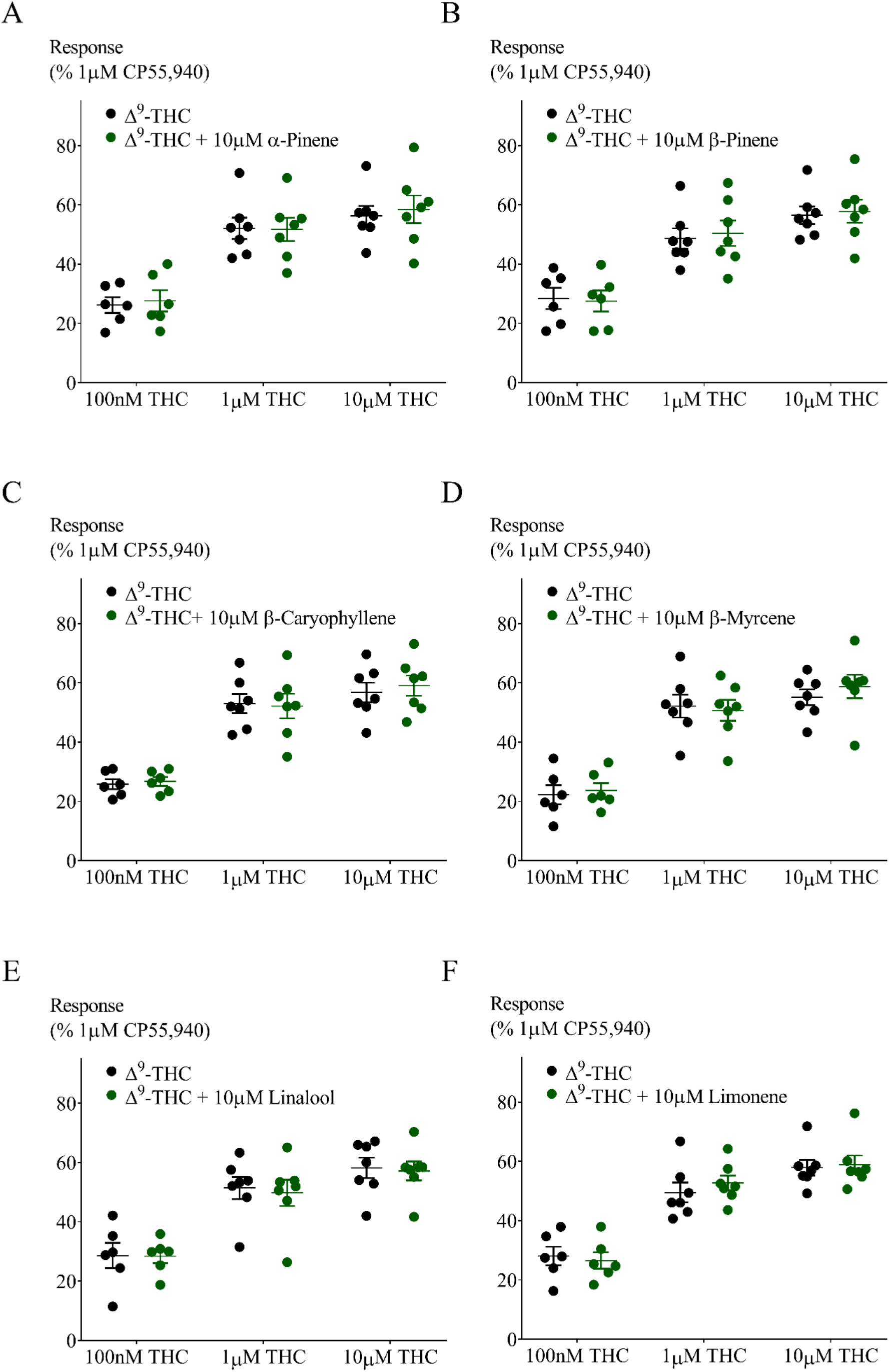
Effect of 10μM terpenoids on Δ^9^-THC induced hyperpolarization in AtT20-CB1. Response to Δ^9^-THC at two sub-maximal and one maximal concentration (n=6-7, SEM, unpaired *t*-test *p >* 0.13). Data presented as % of maximum CP55,940 (1µM) response.

**Fig. 6.**
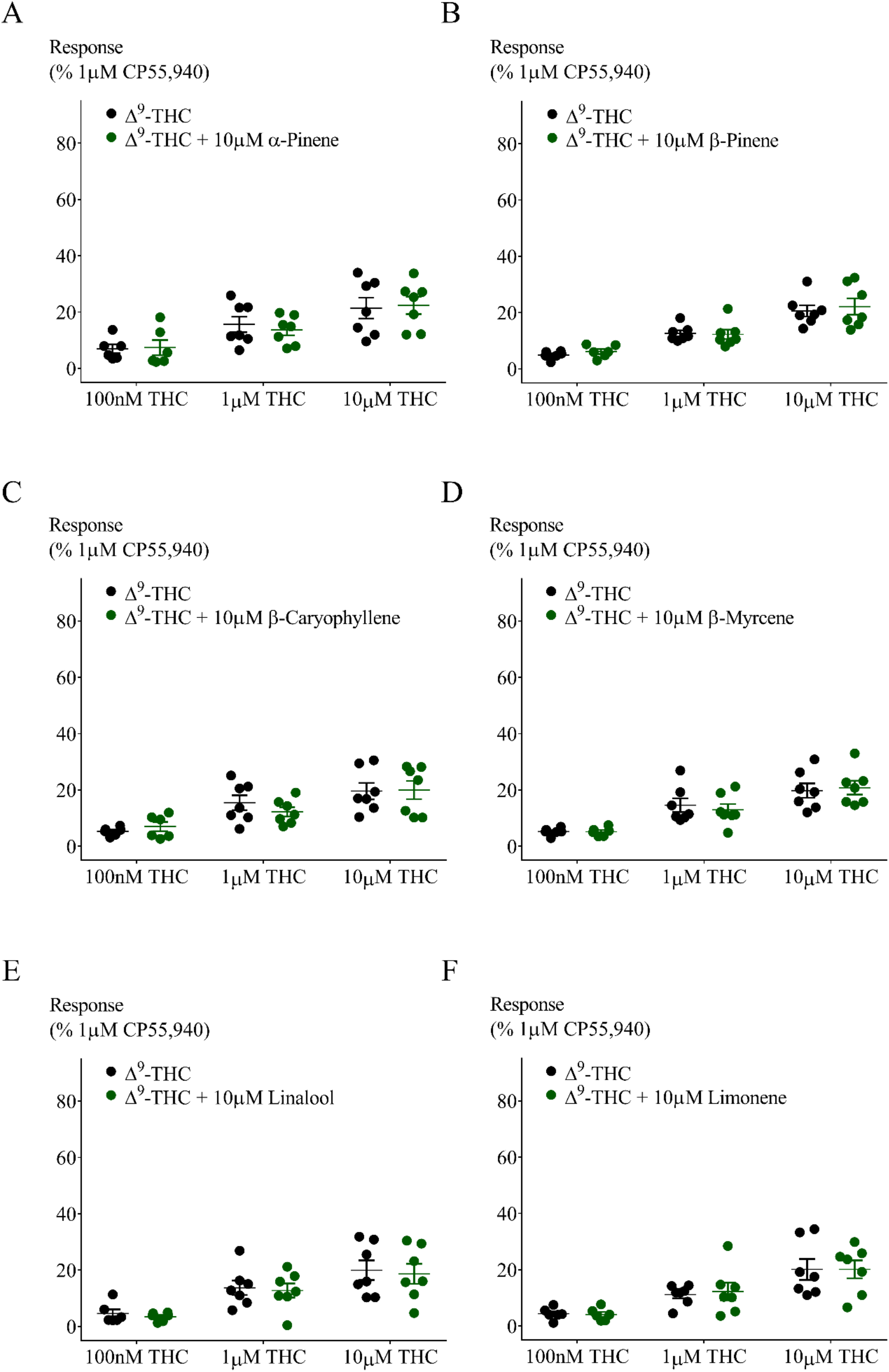
Effect of 10μM terpenoids on Δ^9^-THC induced hyperpolarization in AtT20-CB2. Response to Δ^9^-THC at two sub-maximal and one maximal concentration (n=6-7, SEM, unpaired *t*-test *p >* 0.26). Data presented as % of maximum CP55,940 (1µM) response.

**Fig. 7.**
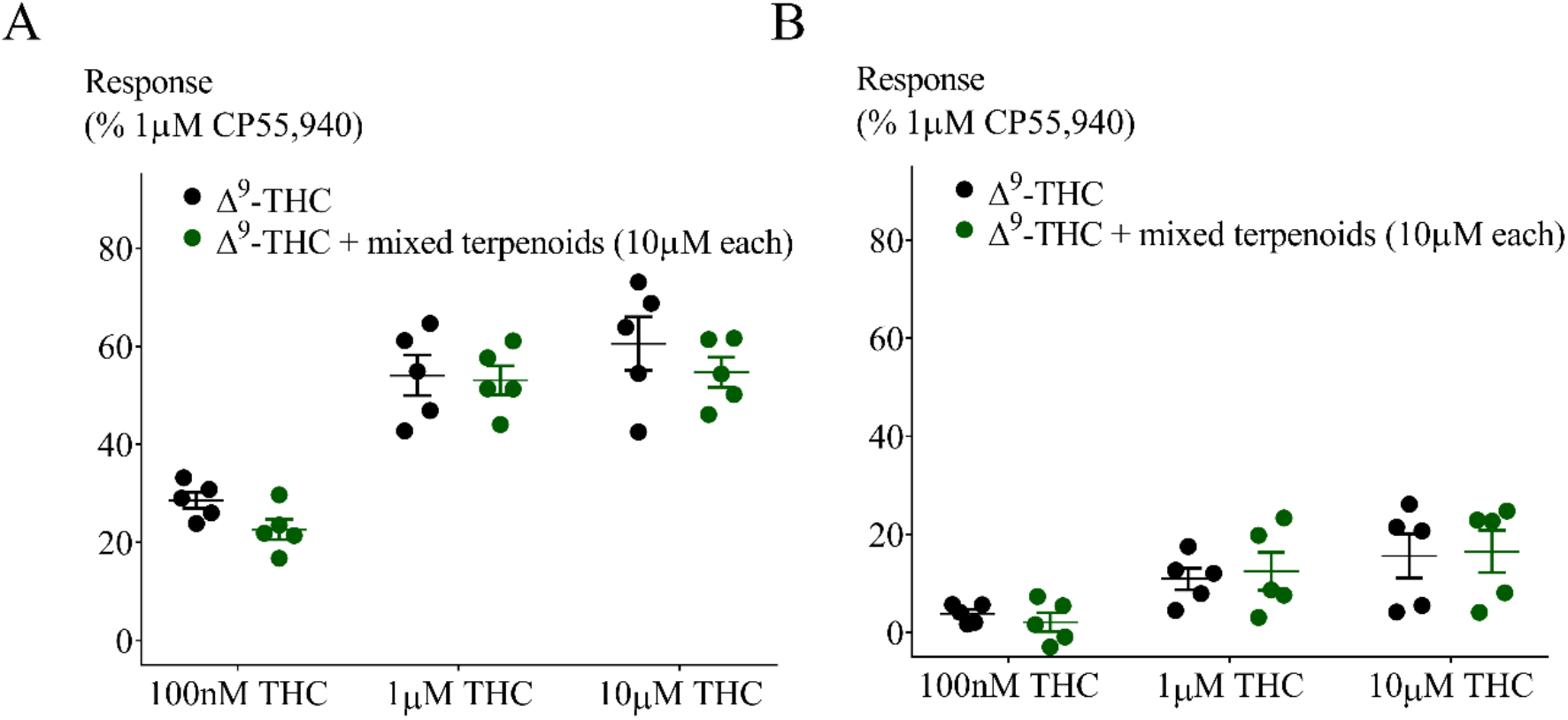
Testing the “Entourage effect”. Effect of combination of six terpenoids at 10μM each on Δ^9^-THC induced hyperpolarization in **A.** AtT20-CB1 and **B.** AtT20-CB2. Response to Δ^9^-THC at two sub-maximal and one maximal concentration (n=5, SEM, unpaired *t*-test *p >* 0.13). Data presented as % of maximum CP55,940 (1µM) response.

### Terpenoids and desensitization in AtT20-CB_1_

We have previously reported desensitization cannabinoid-mediated cellular hyperpolarization in AtT20 cells expressing rat or human CB_1_ receptors ^21, 22^, and we found this reversal of CP55,940-induced hyperpolarization was accelerated by negative allosteric modulators such as ORG27569 and PSNCBAM-1. Therefore, we tested whether terpenoids may act in a similar way to ORG27569 and other negative allosteric modulators, altering desensitization time-course. We used Δ^9^-THC instead of CP55,940, as Δ^9^- THC is the main phytocannabinoid agonist. Prolonged application of Δ^9^-THC (10μM) produced a hyperpolarization that reversed substantially over 30 minutes. Representative traces for this experiment are illustrated in Figure 8A. We measured the peak response to Δ^9^-THC and the signal remaining 30 minutes after agonist exposure and quantified desensitization as a percent decline in the peak response. The Δ^9^-THC (10μM) signal desensitized by 63.3±6.3%, in the presence of the terpenoid mix desensitization was 60.8±4.9% (Fig. 8B). Thus, terpenoids did not interfere with desensitization of CB_1_ signalling produced by Δ^9^-THC. We also assessed the capacity of Δ^9^-THC alone, terpenoids alone (10μM each) or terpenoids combined with Δ^9^-THC to affect somatostatin receptor signalling in AtT20-CB_1_ cells (heterologous desensitization). Somatostatin (100nM) was applied 30 minutes after first drug application (Figs. 8A and 9A and the hyperpolarization produced by somatostatin after Δ^9^-THC, terpenoids alone or Δ^9^-THC with terpenoids were not significantly different to somatostatin alone (*p* > 0.05, Fig. 8B).

**Fig. 8.**
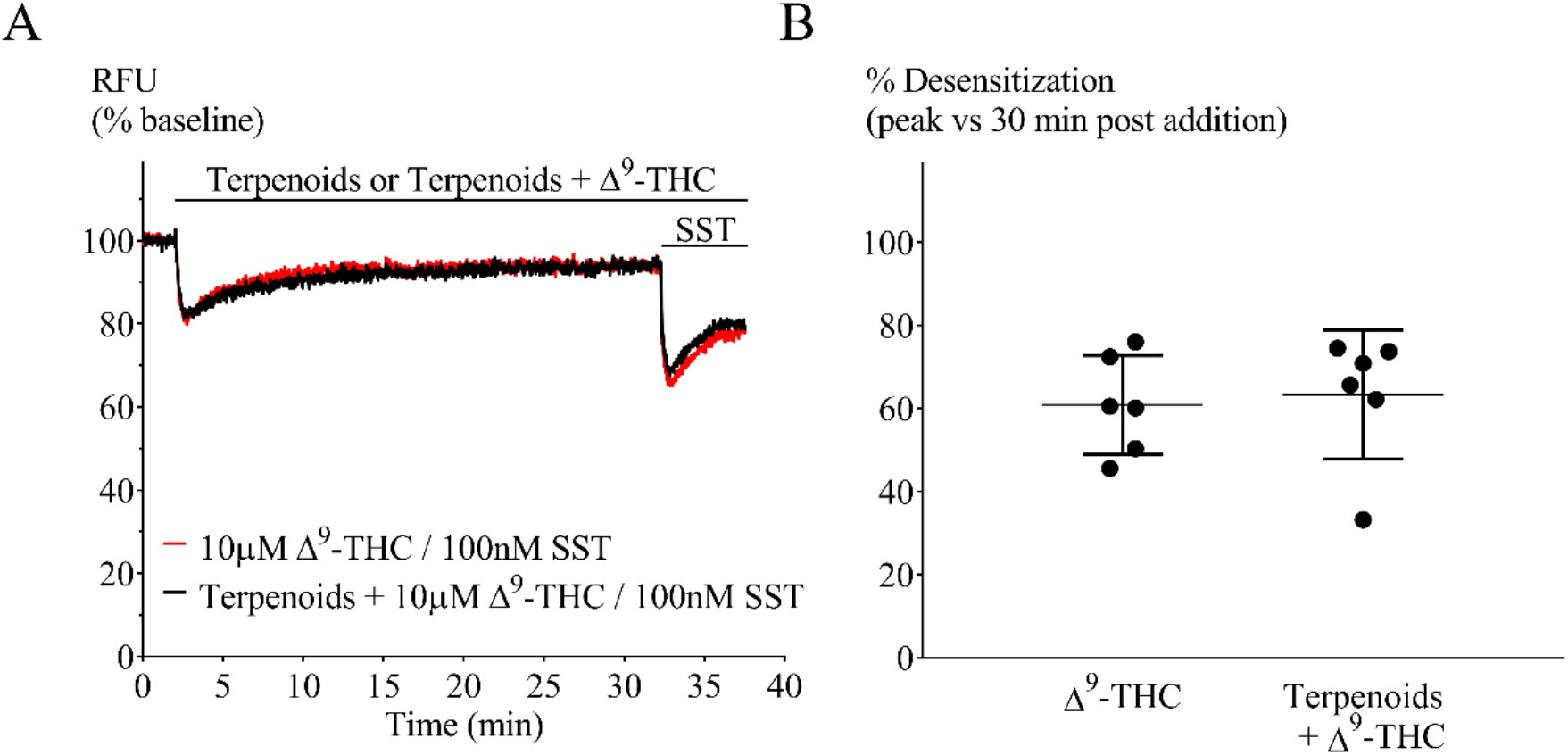
Terpenoids on Δ^9^-THC mediated desensitization in AtT20-CB1. **A.** Representative traces of hyperpolarization and signal desensitization mediated by Δ^9^-THC alone (10µM, black) or with terpenoids (10µM each, red). Cells were then challenged with somatostatin (100nM) after 30 minutes to examine heterologous desensitization. **B.** Percentage desensitization after 30 minutes exposure to Δ^9^-THC alone (10µM) or in the presence of terpenoids (10µM each), compared to peak fluorescence response. Terpenoids did not affect Δ^9^-THC mediated desensitization (n=5, SEM, unpaired *t*-test *p* = 0.76). SST = somatostatin. Drugs were added for the duration of the bar.

**Fig. 9.**
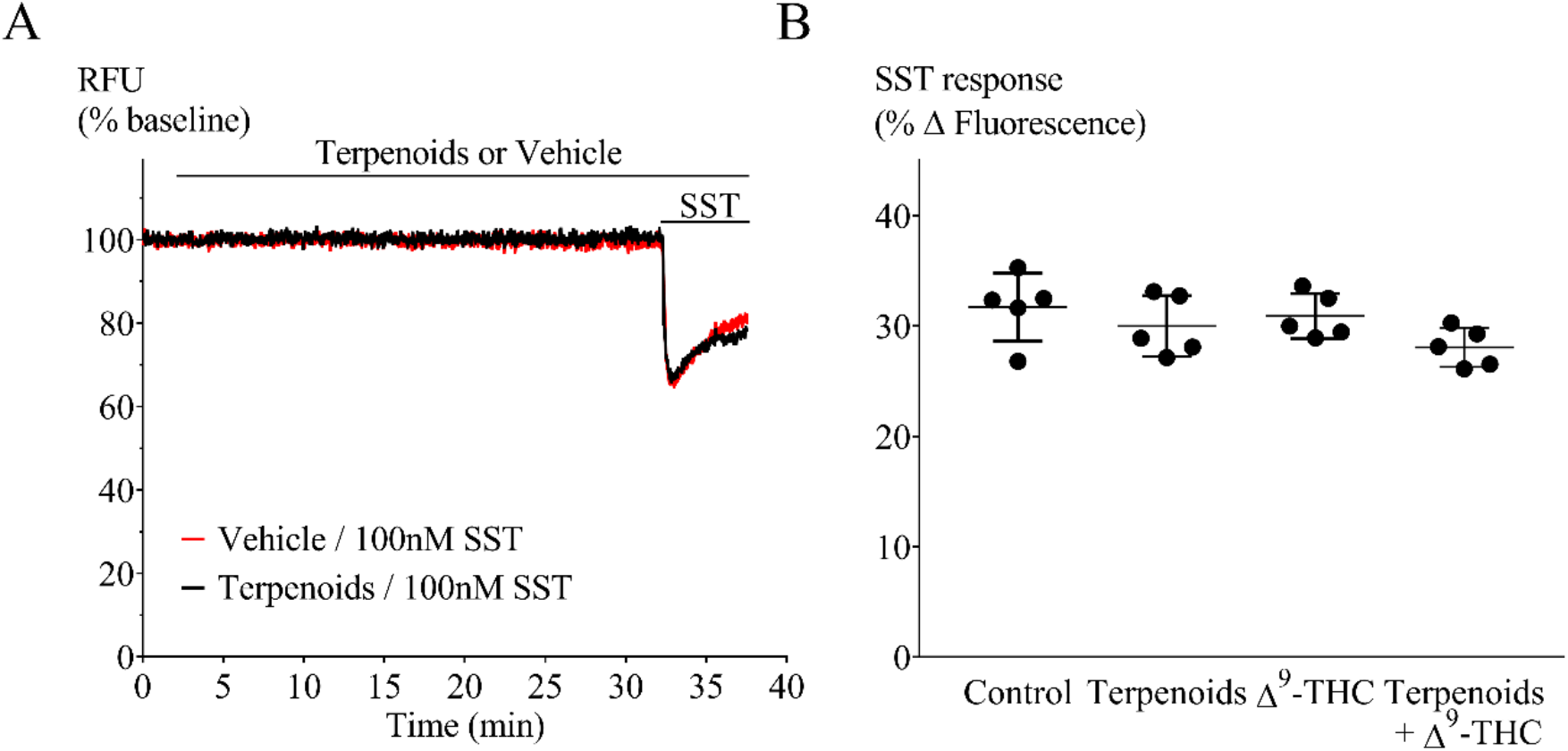
Somatostatin challenge of AtT20-CB1 cells to investigate heterologous desensitization. **A.** Representative traces of cells preincubated with (black) or without (red) terpenoids for 30 minutes before somatostatin (100nM) challenge. **B.** Comparison of peak hyperpolarization (% fluorescence change) obtained after somatostatin (100nM) challenge (n = 5, one-way ANOVA *p >* 0.05). SST = somatostatin. Drugs were added for the duration of the bar.

## Discussion

The principal finding of this study is that agonist activation of CB_1_ and CB_2_ receptors is not obviously altered by any or all of the 6 major terpenoids from *Cannabis sativa*. The terpenoids tested did not activate CB_1_ or CB_2_ by themselves, nor did they modify the signalling of the high efficacy agonist CP55,940 or the lower efficacy agonist Δ^9^-THC. In particular, Δ^9^-THC effects would be expected to be very sensitive to the presence of drugs which inhibited (or enhanced) signalling at the receptor. There are no spare receptors for Δ^9^-THC in this assay, and changes in ligand binding would be directly reflected as a change in the maximum response. The lack of effect of terpenoids on the response to the synthetic cannabinoid CP55,940 indicates that terpenoids do not interfere with maximal cannabinoid receptor-mediated hyperpolarization, suggesting no direct modulation of the potassium channel response. This was confirmed by the lack of effect of terpenoids on the response to somatostatin.

A previous study provided evidence that β-caryophyllene is a CB_2_ agonist^23^. However, we were unable to detect any effect of β-caryophyllene on CB_2_ signalling in the present study. The reasons for this are unclear, but the efficacy of β-caryophyllene has not been defined in cellular assays and may be lower than Δ^9^-THC. The CB_2_ response to even high concentrations of Δ^9^-THC in our assay is small, suggesting that productive coupling of CB_2_ to endogenous potassium channels in AtT-20 cells requires high efficacy agonists. The affinity of β-caryophyllene for CB_2_ (155nM) has been determined in membranes from HEK293 cells heterologously expressing CB_2_^23^, but is not known in intact cells. Its EC_50_ for inhibition of forskolin-induced adenylyl cyclase in CHO-K1 expressing CB_2_ was around 2µM,^23^ suggesting a low functional affinity, which may not be sufficient to significantly affect the rapid response to the higher affinity agonist Δ^9^-THC.

Positive and negative allosteric modulators have been reported for CB_1_^24, 25^, and the effects of several negative allosteric modulators have been defined in the hyperpolarization assay used here.^21^ Both PSNCBAM-1 and ORG27569 enhanced CP55,940 signal desensitization, while PSNCBAM-1 also inhibited the initial CP55,940 hyperpolarization. Co-application of the terpenoids with Δ^9^-THC failed to affect the peak response, or the degree of tachyphylaxis observed over a 30-minute exposure to drug, suggesting that they are not acting as allosteric modulators of this CB_1_ signalling pathway.

A limitation of the present study is that we only examined CB_1_ and CB_2_ signalling through one pathway, involving Gi/o. The hyperpolarization of the AtT20 cells likely represents G-protein mediated activation of inwardly rectifying potassium channels (GIRK), as previously described for CB_1_ and other GPCR in these cells as well as in several different neurons.^26, 27, 28^ Cannabinoid receptors couple to multiple G proteins as well as signalling through other pathways such as those dependent on arrestins and It is possible that entourage effects of terpenoids are mediated through modulation of a subset of the cannabinoid receptor signalling repertoire^26^. CB_1_ and CB_2_ receptors can be activated in a ligand biased manner – the phenomenon where a drug preferentially activates a subset of the signalling pathways that the receptor can access.^29^ In general, this bias has been best defined for G protein coupling versus activation of arrestin-mediated signalling, but to our knowledge there are no examples of cannabinoid ligands only affecting arrestin-mediated signalling.^20, 30^ It remains possible that terpenoids have such an absolute bias, but this would be unprecedented, and in any case recruitment of arrestin would be expected to produce enhanced desensitization of the CB_1_ responses to prolonged agonist exposure ^21, 29^. Any subtle change to receptor signalling should be clear with use of the low efficacy agonist Δ^9^-THC.

Overall, our data suggest that it is unlikely that the terpenoids studied here affect Δ^9^-THC interactions with cannabinoid receptors. However, this is not a definitive rebuttal of the entourage effect. There are many other ways that these molecules could interact with cannabinoids to influence the overall therapeutic and subjective outcomes of cannabis administration and it should be acknowledged that Δ^9^-THC influences signalling at a wide variety of other non-cannabinoid receptor targets (see Banister et al ^31^ for review). These include interaction with metabolic pathways, other G-protein coupled receptors, ligand-gated ion channels, signalling cascades present on the same cells that express cannabinoid receptors, or on other cells up or downstream of the cannabinoid-receptor expressing cells. Terpenoids may even have primary effects on distinct functional modules that together with cannabinoid receptor-modulated pathways are ultimately integrated into a behavioural or physiological output. So the quest for entourage does not end here; in many ways it has only just begun.

---

**Supplementary Fig. S1.**
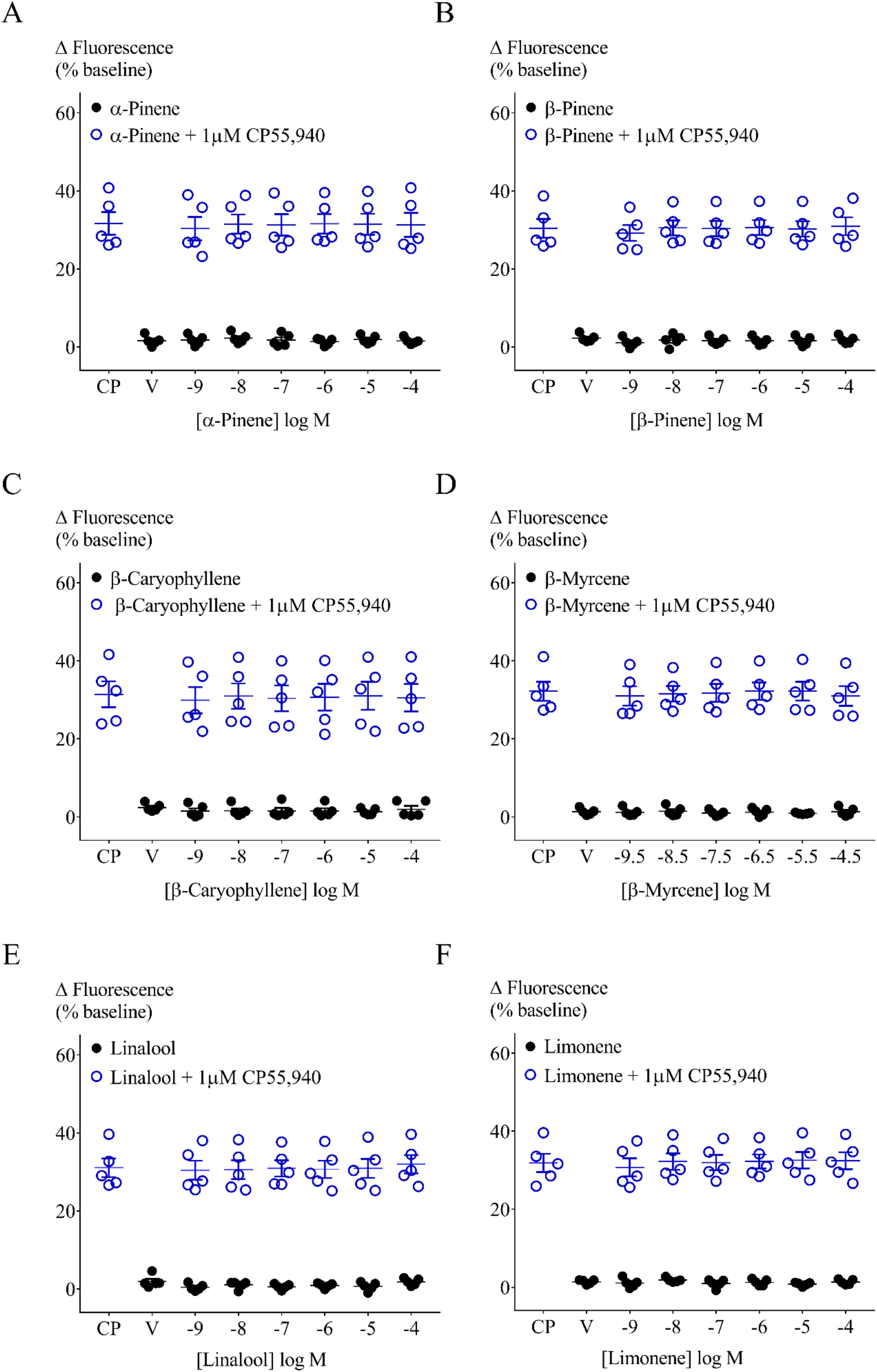
Effect of terpenoids at varying concentrations on AtT20-CB2 membrane potential and on 1µM CP55,940 induced hyperpolarization. Terpenoids (**A.** α-pinene, **B.** β-pinene, **C.** β-caryophyllene, **D.** β-myrcene, **E.** linalool and **F.** Limonene) were added to AtT20-CB2 cells and incubated for 5 minutes. Maximum fluorescence changes were determined and compared to negative control (HBSS, closed circles). No significant fluorescence difference was observed when comparing means of terpenoids and HBSS (n=5, SEM, unpaired *t*-test *p* = 0.72). CP55,940 (1µM) addition to AtT20-CB2 cells induced fluorescence changes from 30.4 ±2.4% to 32.2 ±2.5%. Peak CP55,940 responses were not affected by the presence of terpenoids (open circles, n=5, SEM, unpaired *t*-test *p >* 0.09). V = Vehicle

**Supplementary Fig. S2.**
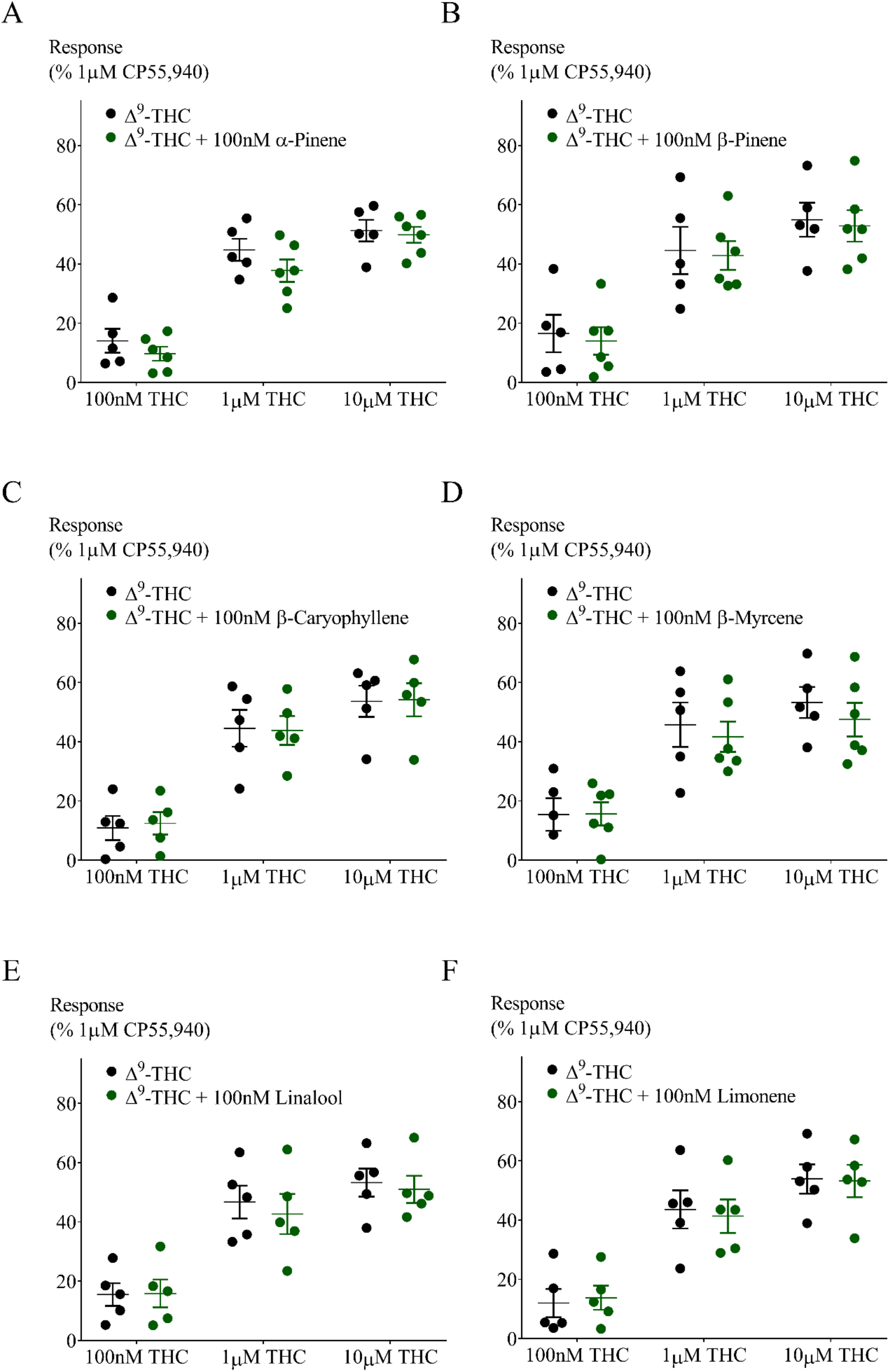
Effect of 100nM Terpenoids on peak hyperpolarization induced by Δ^9^-THC in AtT20- CB1 cells. Response to Δ^9^-THC at two sub-maximal and one maximal concentration (n=5, SEM, unpaired *t*-test *p >* 0.24). Data presented as % of maximum CP55,940 (1µM) response.

**Supplementary Fig. S3.**
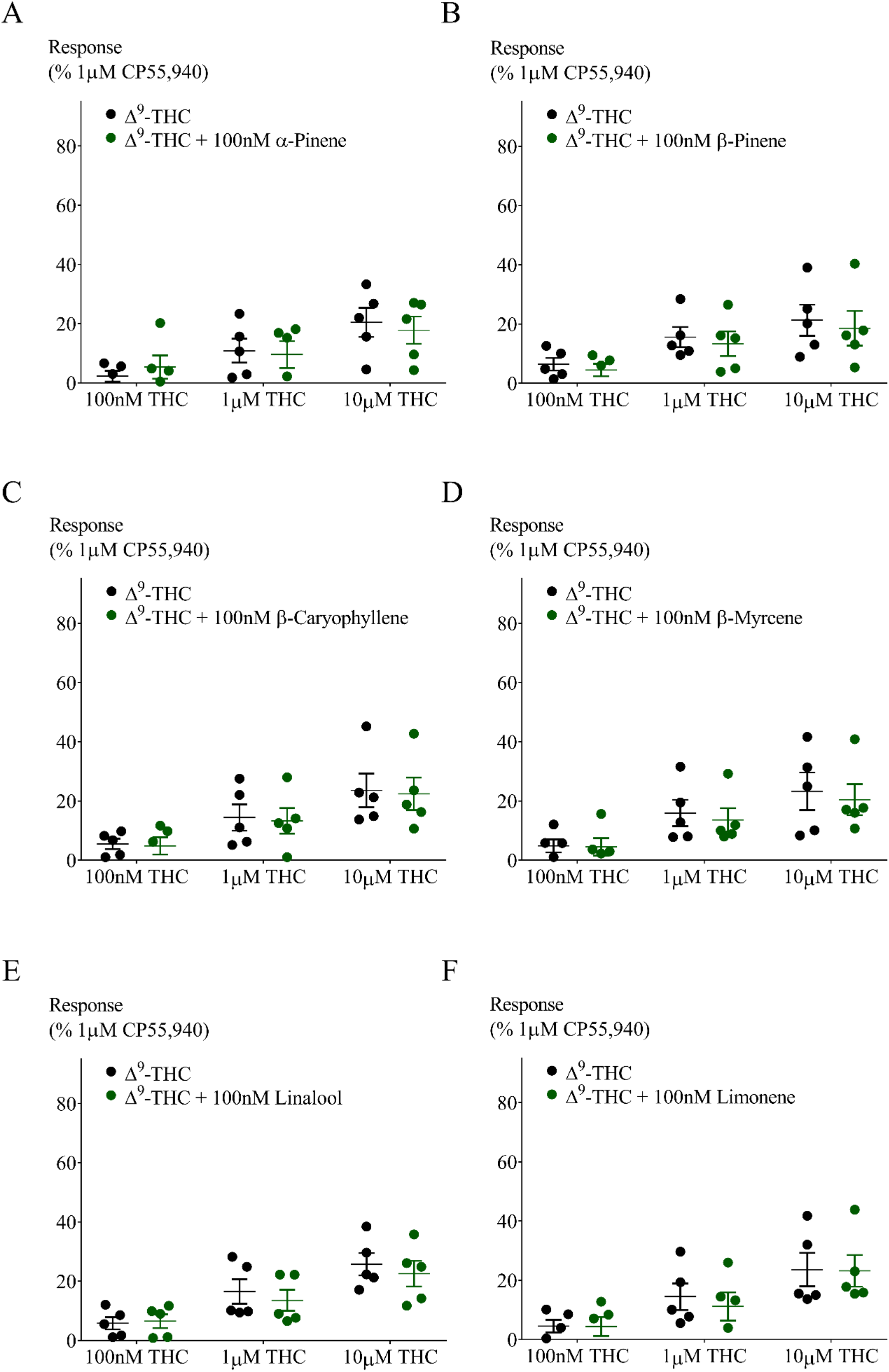
Effect of 100nM Terpenoids on peak hyperpolarization induced by Δ^9^-THC in AtT20- CB2 cells. Response to Δ^9^-THC at two sub-maximal and one maximal concentration (n=5, SEM, unpaired *t*-test *p >* 0.50). Data presented as % of maximum CP55,940 (1µM) response.

## Abbreviations

β-Car: β-caryophyllene
CBD: cannabidiol
HBSS: Hank’s balanced salt solution
RFU: relative fluorescence units
SEM: standard error of the mean
SST: somatostatin
Δ^9^-THC: Δ^9^-tetrahydrocannabinol

## References

1. Russo EB. Taming THC: potential cannabis synergy and phytocannabinoid-terpenoid entourage effects. Br J Pharmacol. 2011;163(7):1344–1364.

2. Hanuš LO, Meyer SM, Muñoz E, et al. Phytocannabinoids: a unified critical inventory. Natural Product Reports. 2016;33(12):1357–1392.

3. Ibrahim EA, Wang M, Radwan MM, et al. Analysis of Terpenes in Cannabis sativa L. Using GC/MS: Method Development, Validation, and Application. Planta medica. 2019.

4. Pertwee RG. Handbook of cannabis. 1st ed. Oxford, United Kingdom; New York, NY: Oxford University Press; 2014.

5. Sexton M, Shelton K, Haley P, et al. Evaluation of Cannabinoid and Terpenoid Content: Cannabis Flower Compared to Supercritical CO2 Concentrate. Planta medica. 2018;84(4):234–241.

6. Casey SL, Atwal N, Vaughan CW. Cannabis constituent synergy in a mouse neuropathic pain model. Pain. 2017;158(12):2452–2460.

7. Englund A, Freeman TP, Murray RM, et al. Can we make cannabis safer? Lancet Psychiatry. 2017;4(8):643–648.

8. Russo E, Guy GW. A tale of two cannabinoids: the therapeutic rationale for combining tetrahydrocannabinol and cannabidiol. Med Hypotheses. 2006;66(2):234–246.

9. Haney M, Malcolm RJ, Babalonis S, et al. Oral Cannabidiol does not Alter the Subjective, Reinforcing or Cardiovascular Effects of Smoked Cannabis. Neuropsychopharmacology. 2016;41(8):1974–1982.

10. Ilan AB, Gevins A, Coleman M, et al. Neurophysiological and subjective profile of marijuana with varying concentrations of cannabinoids. Behav Pharmacol. 2005;16(5-6):487–496.

11. Laprairie RB, Bagher AM, Kelly ME, et al. Cannabidiol is a negative allosteric modulator of the cannabinoid CB1 receptor. Br J Pharmacol. 2015;172(20):4790–4805.

12. Russo EB. The Case for the Entourage Effect and Conventional Breeding of Clinical Cannabis: No “Strain,” No Gain. Frontiers in plant science. 2018;9:1969.

13. Lewis MA, Russo EB, Smith KM. Pharmacological Foundations of Cannabis Chemovars. Planta medica. 2018;84(04):225–233.

14. Russo EB, Marcu J. Cannabis Pharmacology: The Usual Suspects and a Few Promising Leads. Adv Pharmacol. 2017;80:67–134.

15. Potter D. The propagation, characterisation and optimisation of Cannabis sativa L. as a phytopharmaceutical [Doctoral Thesis]. London: King’s College; 2009.

16. Suraev A, Lintzeris N, Stuart J, et al. Composition and Use of Cannabis Extracts for Childhood Epilepsy in the Australian Community. Scientific reports. 2018;8(1):10154.

17. Banister SD, Longworth M, Kevin R, et al. Pharmacology of Valinate and tert-Leucinate Synthetic Cannabinoids 5F-AMBICA, 5F-AMB, 5F-ADB, AMB-FUBINACA, MDMB-FUBINACA, MDMB-CHMICA, and Their Analogues. ACS Chem Neurosci. 2016;7(9):1241–1254.

18. Knapman A, Santiago M, Du YP, et al. A continuous, fluorescence-based assay of mu-opioid receptor activation in AtT-20 cells. J Biomol Screen. 2013;18(3):269–276.

19. Gunther T, Culler M, Schulz S. Research Resource: Real-Time Analysis of Somatostatin and Dopamine Receptor Signaling in Pituitary Cells Using a Fluorescence-Based Membrane Potential Assay. Mol Endocrinol. 2016;30(4):479–490.

20. Soethoudt M, Grether U, Fingerle J, et al. Cannabinoid CB2 receptor ligand profiling reveals biased signalling and off-target activity. Nat Commun. 2017;8:13958.

21. Cawston EE, Redmond WJ, Breen CM, et al. Real-time characterization of cannabinoid receptor 1 (CB1) allosteric modulators reveals novel mechanism of action. Br J Pharmacol. 2013;170(4):893–907.

22. Cawston EE, Connor M, Di Marzo V, et al. Distinct Temporal Fingerprint for Cyclic Adenosine Monophosphate (cAMP) Signaling of Indole-2-carboxamides as Allosteric Modulators of the Cannabinoid Receptors. Journal of medicinal chemistry. 2015;58(15):5979–5988.

23. Gertsch J, Leonti M, Raduner S, et al. Beta-caryophyllene is a dietary cannabinoid. Proc Natl Acad Sci U S A. 2008;105(26):9099–9104.

24. Price MR, Baillie GL, Thomas A, et al. Allosteric modulation of the cannabinoid CB1 receptor. Mol Pharmacol. 2005;68(5):1484–1495.

25. Ignatowska-Jankowska BM, Baillie GL, Kinsey S, et al. A Cannabinoid CB1 Receptor-Positive Allosteric Modulator Reduces Neuropathic Pain in the Mouse with No Psychoactive Effects. Neuropsychopharmacology. 2015;40(13):2948–2959.

26. Bacci A, Huguenard JR, Prince DA. Long-lasting self-inhibition of neocortical interneurons mediated by endocannabinoids. Nature. 2004;431(7006):312–316.

27. Marinelli S, Pacioni S, Cannich A, et al. Self-modulation of neocortical pyramidal neurons by endocannabinoids. Nature neuroscience. 2009;12(12):1488–1490.

28. Felder CC, Joyce KE, Briley EM, et al. Comparison of the pharmacology and signal transduction of the human cannabinoid CB1 and CB2 receptors. Mol Pharmacol. 1995;48(3):443–450.

29. Ibsen MS, Connor M, Glass M. Cannabinoid CB1 and CB2 Receptor Signaling and Bias. Cannabis Cannabinoid Res. 2017;2(1):48–60.

30. Atwood BK, Wager-Miller J, Haskins C, et al. Functional selectivity in CB(2) cannabinoid receptor signaling and regulation: implications for the therapeutic potential of CB(2) ligands. Mol Pharmacol. 2012;81(2):250–263.

31. Banister SD, Arnold JC, Connor M, et al. Dark Classics in Chemical Neuroscience: Δ9-Tetrahydrocannabinol. ACS Chemical Neuroscience. 2019.

